# GHSR-1a is not Required for Ghrelin’s Anti-inflammatory and Fat-sparing Effects in Cancer Cachexia

**DOI:** 10.1101/866376

**Authors:** Haiming Liu, Jiaohua Luo, Bobby Guillory, Ji-an Chen, Pu Zang, Jordan K. Yoeli, Yamileth Hernandez, Ian (In-gi) Lee, Barbara Anderson, Mackenzie Storie, Alison Tewnion, Jose M. Garcia

**Affiliations:** Geriatric Research, Education and Clinical Center, Veterans Affairs Puget Sound Health Care System, Seattle, 98108 WA, USA; Gerontology and Geriatric Medicine, University of Washington Department of Medicine, Seattle, 98195 WA, USA; Division of Endocrinology, Diabetes and Metabolism, MCL, Center for Translational Research on Inflammatory Diseases, Michael E. DeBakey Veterans Affairs Medical Center, Dept. of Medicine, Baylor College of Medicine, Houston, TX 77030, USA; Department of Environmental Hygiene, College of Preventive Medicine, Army Medical University, Chongqing, 400038, China; Department of Health Education, College of Preventive Medicine, Army Medical University, Chongqing, 400038, China; Department of Endocrinology, Nanjing Jinling Hospital, Nanjing, 210002, China

**Keywords:** cachexia, cancer, muscle, ghrelin, adipose tissue

## Abstract

Adipose tissue (AT) atrophy is a hallmark of cancer cachexia contributing to increased morbidity/mortality. Ghrelin has been proposed as a treatment for cancer cachexia partly by preventing AT atrophy. However, the mechanisms mediating ghrelin’s effects are incompletely understood, including the extent to which its only known receptor, GHSR-1a, is required for these effects. This study characterizes the pathways involved in AT atrophy in the Lewis Lung Carcinoma (LLC)-induced cachexia model and those mediating the effects of ghrelin in *Ghsr^+/+^* and *Ghsr^−/−^* mice. We show that LLC causes AT atrophy by inducing anorexia, and increasing AT inflammation, thermogenesis and energy expenditure. These changes were greater in *Ghsr^−/−^*. Ghrelin administration prevented LLC-induced anorexia only in *Ghsr^+/+^*, but prevented WAT inflammation and atrophy in both genotypes, although its effects were greater in *Ghsr^+/+^*. LLC-induced increases in BAT inflammation, WAT and BAT thermogenesis, and energy expenditure were not affected by ghrelin. In conclusion, ghrelin ameliorates WAT inflammation, fat atrophy and anorexia in LLC-induced cachexia. GHSR-1a is required for ghrelin’s orexigenic effect but not for its anti-inflammatory or fat-sparing effects.

## INTRODUCTION

Every year, over 1,500,000 individuals in the US are diagnosed with cancer. Cachexia (involuntary loss of muscle and adipose tissue) is present in up to 80% of cancer patients, is strongly associated with higher morbidity and mortality, and is reported as the direct cause of death in 20-40% of these patients (Dewys, Begg et al., 1980, Fearon, Strasser et al., 2011). Adipose tissue, once considered only a high-energy fuel reserve, has emerged recently as an active metabolic organ modulating inflammation, energy expenditure and food intake in non-cancer settings (You & Nicklas, 2006). Accelerated loss of adipose tissue plays an important role in cancer cachexia contributing significantly to the increased morbidity and mortality seen in this setting (Fouladiun, Korner et al., 2005).

Increased inflammation is common in the setting of cancer (Garcia, Garcia-Touza et al., 2005) and is associated with adipose tissue wasting in human studies (Lerner, Hayes et al., 2015). White adipose tissue (WAT) is a significant source of inflammatory cytokines accounting for more than 30% of circulating interleukin (IL)-6 (Michaud, Boulet et al., 2014) and this and other inflammatory cytokines have been linked to WAT atrophy in the setting of cancer (Petruzzelli, Schweiger et al., 2014, Tsoli & Robertson, 2013, Tsoli, Swarbrick et al., 2016). Also, a phenotypic switch from WAT to brown adipose tissue (BAT) known as “browning” is thought to contribute to the overall increase in energy expenditure and WAT atrophy seen in cancer cachexia (Petruzzelli et al., 2014). Nevertheless, the mechanisms regulating adipose tissue atrophy and dysfunction in this setting are incompletely understood.

Ghrelin, originally identified as the endogenous ligand for the growth hormone secretagogue receptor (GHSR)-1a, has emerged as a pleiotropic hormone that regulates body weight, body composition and energy expenditure (Muller & Tschop, 2013). In non-cancer models, it has been shown to increase food intake by activating neuropeptide Y and agouti-related peptide-secreting neurons in the hypothalamus and to have direct effects on adipocytes (Kos, Harte et al., 2009, Muller & Tschop, 2013, Perez-Tilve, Heppner et al., 2011). Ghrelin has also been proposed as a promising target for cancer cachexia and it has been shown to prevent fat atrophy in tumor-bearing animals and in patients with cancer cachexia (Chen, Splenser et al., 2015, Garcia, Boccia et al., 2015, Garcia, Scherer et al., 2013b). However, the mechanisms mediating these effects are incompletely understood. Interestingly, emerging data suggest that some of these effects are independent of the only ghrelin receptor identified to date, GHSR-1a (Kojima, Hosoda et al., 1999, Smith, Van der Ploeg et al., 1997).

The objectives of this study were to characterize the pathways involved in adipose tissue atrophy in the Lewis Lung Carcinoma (LLC)-induced cachexia model and to determine the pathways mediating the effects of ghrelin on adipose tissue in this setting, including the relative contribution of GHSR-1a.

## RESULTS

We utilized C57/BL6 congenic mice with (*Ghsr^+/+^*) or without GHSR-1a (*Ghsr*^−/−^). Five to seven-month-old male *Ghsr^+/+^* and *Ghsr^−/−^* mice were inoculated with 1×10^6^ heat-killed (HK, control) or live LLC cells in the right flank. When the tumor was palpable (approximately 1 wk after implantation), tumor-bearing mice were injected with vehicle (saline solution, tumor-vehicle, TV) or ghrelin (0.8 mg/kg, tumor-ghrelin, TG) subcutaneously (s.q.) twice/day, while HK mice were injected with vehicle until the end of the experiments (2 weeks after the tumor became palpable). Body weight and fat mass were measured by nuclear magnetic resonance (NMR) before tumor implantation and 2 weeks after tumors were noted. Brown adipose tissue (BAT) and inguinal and epididymal white adipose tissue (iWAT, eWAT) were collected and weighed upon sacrificing animals 2 weeks after tumors were noted. We confirmed that *Ghsr^−/−^* mice did not express *Ghsr* globally by genotyping. Also, there was no expression of *Ghsr* in neither iWAT or BAT on either genotype (Supplemental Fig.1).

### Ghrelin prevents tumor-induced weight loss and adipose tissue atrophy only partially via GHSR-1a

LLC tumor implantation induced significant decreases in carcass weight in both genotypes; although, the decrease was more profound in *Ghsr^−/−^* than in *Ghsr^+/+^* mice (Fig. 1A, genotype effect: *p* < 0.001). The same pattern was seen in whole body fat mass measured by NMR (Fig. 1B, genotype effect: *p* = 0.002) as well as in iWAT and eWAT pad weights measured upon dissection (Fig. 1C, genotype effect on iWAT: *p* = 0.043). These changes were fully prevented by ghrelin administration in *Ghsr^+/+^* tumor-bearing animals and partially prevented in *Ghsr^−/−^* animals. *Ghsr^−/−^* mice exhibited significantly less food intake *versus Ghsr^+/+^* mice during daytime (genotype effect: *p* = 0.018) and tumor-bearing mice showed less food intake than controls, although this difference only reached significant for the TG group at nighttime (Figure 1D). LLC-induced decreases in food intake were prevented by ghrelin during daytime (6am – 6pm) only in *Ghsr^+/+^*.

**Figure 1.**
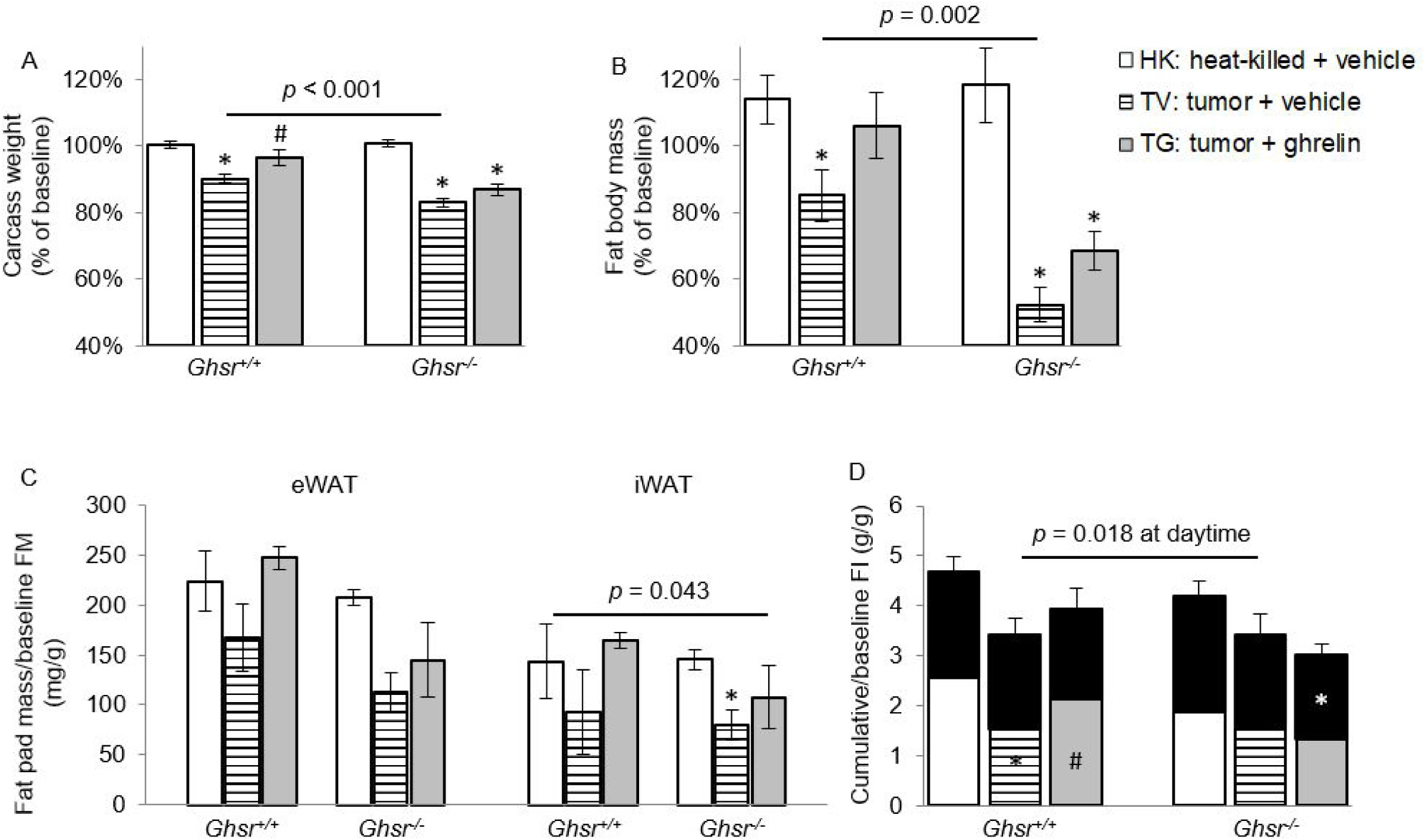
Effects of ghrelin on body weight, fat mass, and food intake in LLC-induced cachexia. HK: heat-killed + vehicle; TV: tumor + vehicle; TG: tumor + ghrelin. Changes in (A) body weight (carcass weight, n = 8-10) and (B) fat body mass by NMR expressed as % change from baseline (n = 8-10). (C) Fat pad mass normalized to baseline NMR fat mass (mg/g, n = 4-6). (D) Average cumulative food intake (FI) normalized to baseline FI (g/g, black areas represent food intake in the nighttime, and the bottom areas in the bars represent food intake in the daytime, n = 4-6). * *p* < 0.05 compared to HK within the same genotype. # *p* <0.05 compared to TV within the same genotype. In panel D, differences in daytime are shown at the lower part of the bars; differences in nighttime are shown at the upper part of the bars. Genotype effects are shown in *p*-values above the corresponding figures (*p* < 0.05). Data are shown as mean ± SE.

### Ghrelin attenuates tumor-induced inflammation in iWAT but not in iBAT or in circulation

In *Ghsr^+/+^* animals, protein level for the pro-inflammatory cytokines IL-1β and TNF in iWAT were increased in tumor-bearing mice and ghrelin prevented these increases (Fig 2A, C). IL-6 and the macrophage marker monocyte chemoattractant protein-1 (MCP-1), a key chemokine responsible for migration and infiltration of monocytes/macrophages (Deshmane, Kremlev et al., 2009), followed a similar pattern although the differences did not reach statistical significance (Fig. 2B, D). Interestingly, in *Ghsr^−/−^* mice LLC-induced IL-6 level increases in iWAT appear to be dampened; whereas, MCP-1 levels were not affected by LLC or by ghrelin. Immunohistochemistry staining shows complete co-localization of IL-6 and TNF with F4/80, a marker of macrophages in mice, demonstrating that the source of these cytokines in iWAT are macrophages (Fig 2 E-F). High resolution images of immunohistochemistry staining in iWAT are demonstrated in Supplemental Fig. 2.

**Figure 2.**
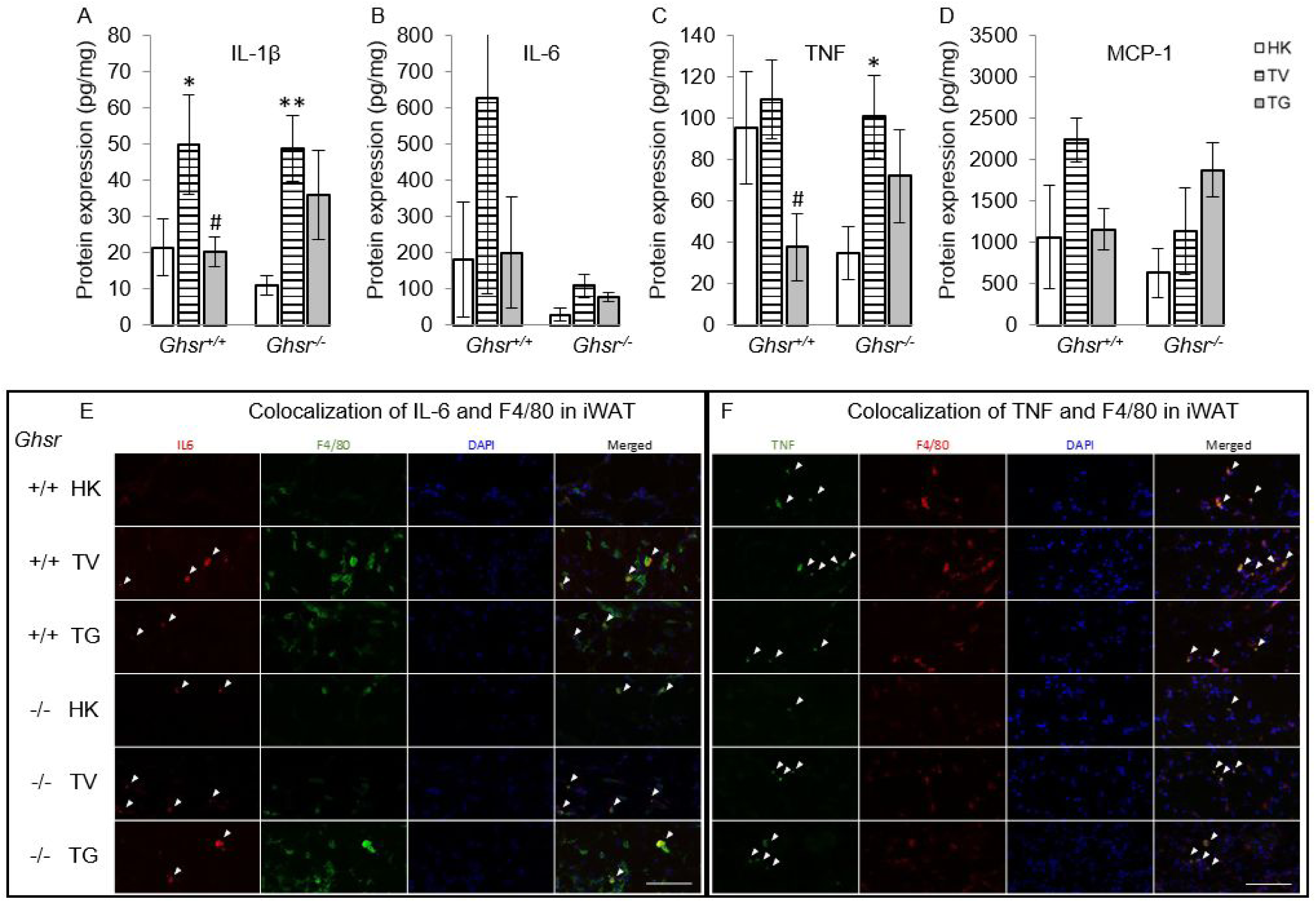
Effects of ghrelin on LLC-induced changes in inflammation and macrophages in iWAT. HK: heat-killed + vehicle; TV: tumor + vehicle; TG: tumor + ghrelin. Protein levels of inflammatory markers (A)IL-1β, (B) IL-6, and (C) TNF; and (D) macrophage marker MCP-1 in iWAT (pg/mg). \**p* < 0.05; \*\**p* < 0.01 compared to HK within the same genotype. # *p* < 0.05 compared to TV within the same genotype. No genotype difference was detected. Data are shown as mean ± SE. n = 6-7/group. (E-F) Colocalization of inflammation and macrophages in iWAT. (E) Representative images of colocalization of inflammatory marker IL-6 and macrophage marker F4/80 in iWAT (IL-6 in Texas red; F4/80 in FITC green; nuclei in DAPI blue). (F) Representative images of colocalization of inflammatory marker TNF and macrophage marker F4/80 in iWAT (TNF in FITC green; F4/80 in Texas red; nuclei in DAPI blue). Positively stained inflammatory markers and colocalizations with macrophages are indicated by the white arrows. Scale bars, 100 µm.

In BAT, all the inflammatory markers were generally lower than in WAT. IL-1β was increased in both genotypes (Fig. 3A) and MCP-1 only in *Ghsr^−/−^* (Fig. 3D). Ghrelin did not significantly affect these changes. IL-6 and TNF levels were not significantly different between groups (Fig. 3B-C). Nevertheless, immunohistochemistry analysis shows similar results as in iWAT suggesting that IL-6 and TNF in BAT were also derived exclusively from macrophages (Fig. 3 E-F). High resolution images of immunohistochemistry staining in BAT are demonstrated in Supplemental Fig. 3. Plasma cytokine and MCP-1 levels followed a different pattern than those seen in adipose tissue being increased by LLC and not modified by ghrelin (Supplemental Fig. 4).

**Figure 3.**
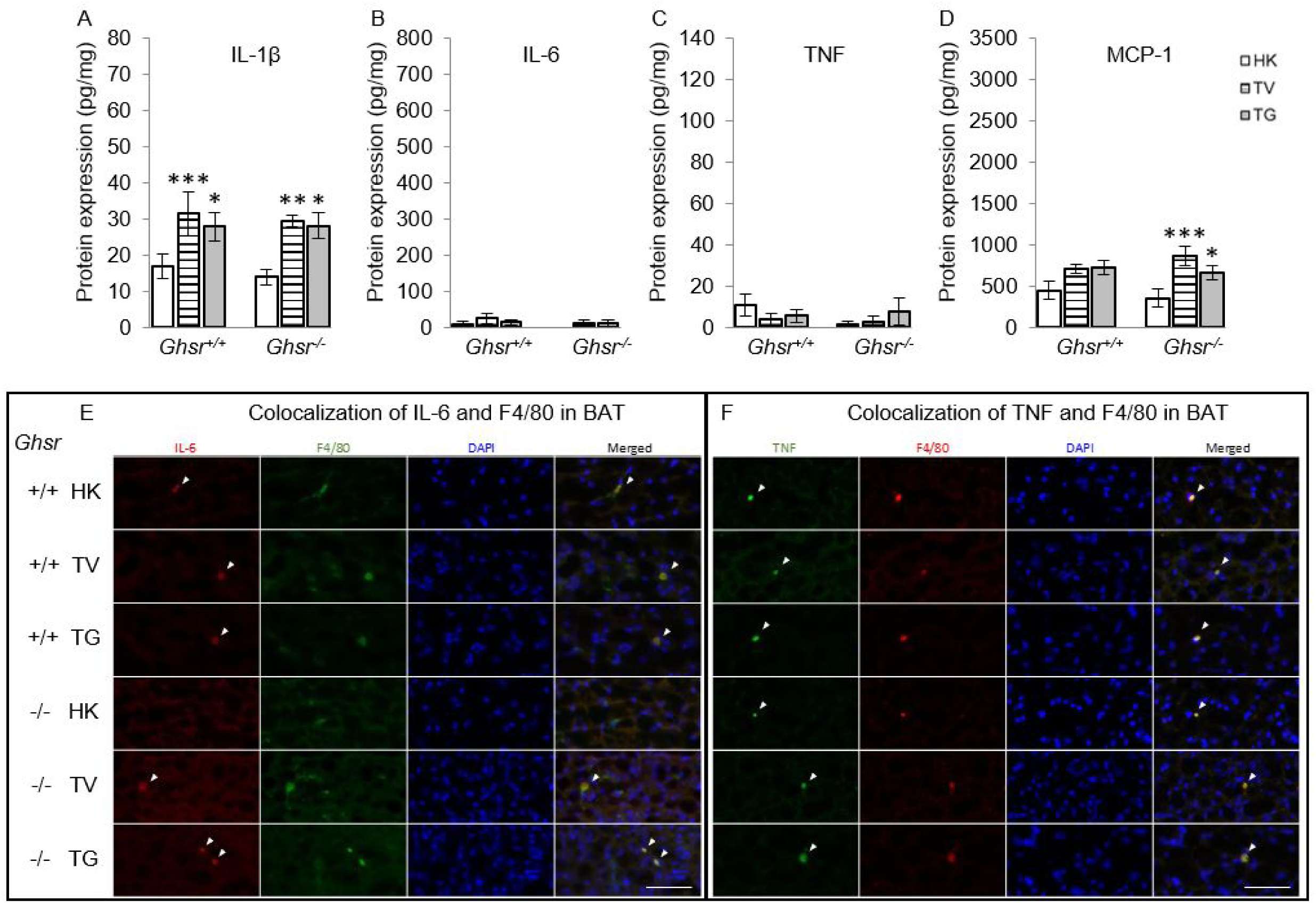
Effects of ghrelin on LLC-induced changes in inflammation and macrophages in BAT. HK: heat-killed + vehicle; TV: tumor + vehicle; TG: tumor + ghrelin. Protein levels of inflammatory markers (A)IL-1β, (B) IL-6, and (C) TNF; and (D) macrophage marker MCP-1 in iWAT (pg/mg). * *p* < 718 0.05; ** *p* < 0.01; *** *p* < 0.001 compared to HK within the same genotype. # *p* < 0.05; ### *p* < 0.001 compared to TV within the same genotype. No genotype difference was detected. Data are shown as mean ± SE. n = 6-7/group. (E-F) Colocalization of inflammation and macrophages in BAT. (E) Representative images of colocalization of inflammatory marker IL-6 and macrophage marker F4/80 in BAT (IL-6 in Texas red; F4/80 in FITC green; nuclei in DAPI blue). (F) Representative images of colocalization of inflammatory marker TNF and macrophage marker F4/80 in BAT (TNF in FITC green; F4/80 in Texas red; nuclei in DAPI blue). Positively stained inflammatory markers and colocalizations with macrophages are indicated by the white arrows. Scale bars, 100 µm.

### Ghrelin does not prevent the increases in UCP-1 induced by LLC in iWAT or BAT

Thermogenesis in BAT is activated by uncoupling protein-1 (UCP-1) by de-coupling oxidative phosphorylation from ATP synthesis and dissipating heat in the inner mitochondrial membrane (Puigserver, Wu et al., 1998). A similar process has been reported in WAT which has been described as “fat browning” with transformation of “white” to “beige” adipocytes (Rosen & Spiegelman, 2014, Wu, Bostrom et al., 2012). To test the effect of LLC and the role of ghrelin and GHSR-1a on this pathway, we quantified UCP-1 levels in iWAT and BAT using immunohistochemistry (IHC) by normalizing the positively-stained area to the total cross-sectional area of the adipose tissue. Tumor implantation induced increases in UCP-1 expression in iWAT and BAT in both genotypes and these increases were more pronounced in *Ghsr^−/−^* than in *Ghsr^+/+^* (Fig 4 A-D, genotype effect in BAT: *p* = 0.005). In iWAT, the LLC-induced UCP-1 increase only reached significance in the tumor-bearing *Ghsr^−/−^* mice and no significant effect of ghrelin was observed. In BAT, the positively stained UCP-1 area increased with tumor implantation from 22% to 59% in *Ghsr^+/+^* and from 35% to 70% in *Ghsr^−/−^* mice. However, no effect of ghrelin on reducing UCP-1 in BAT was observed.

**Figure 4.**
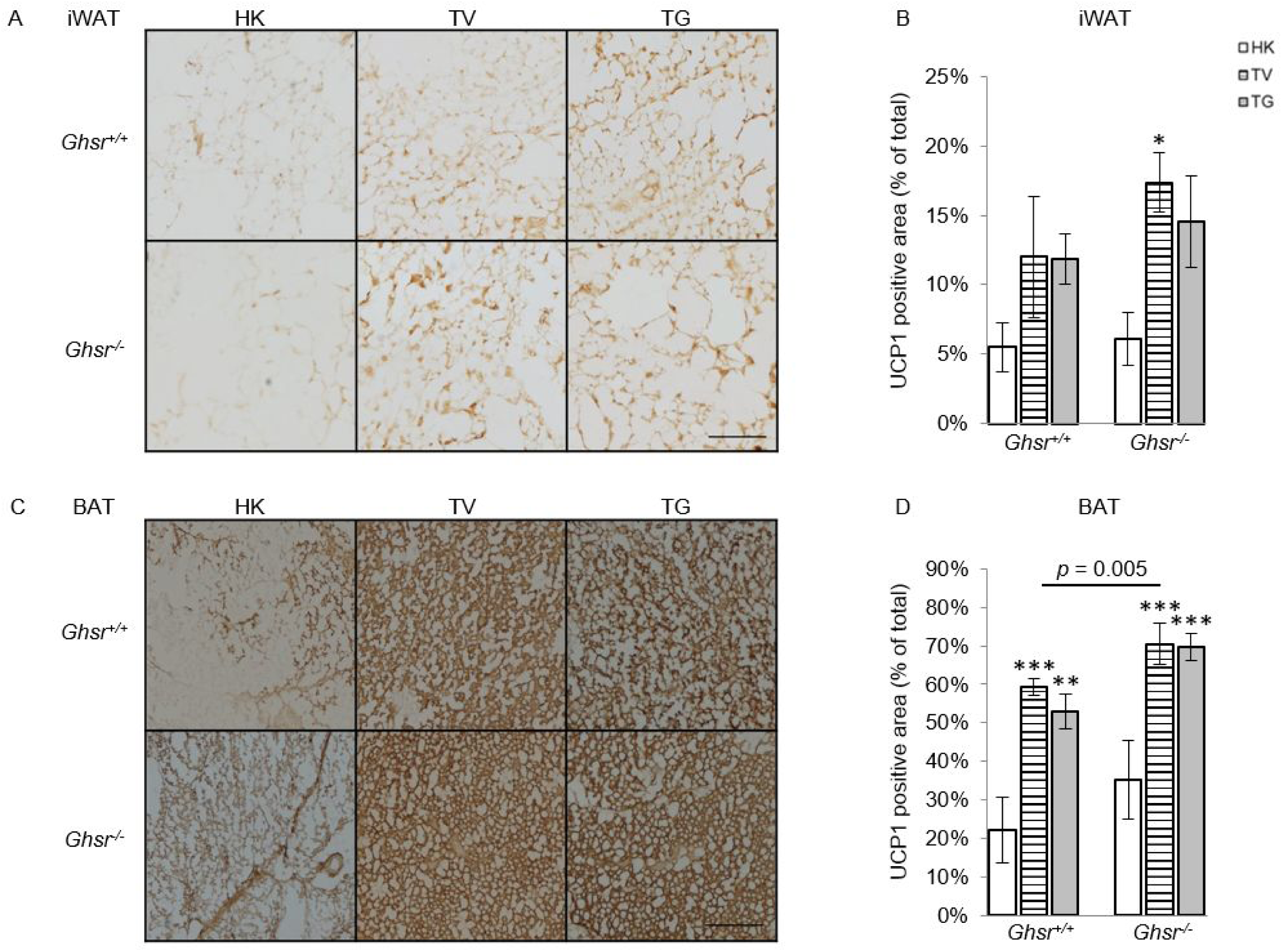
Expression of UCP-1 in iWAT and BAT. HK: heat-killed + vehicle; TV: tumor + vehicle; TG: tumor + ghrelin. (A) Representative IHC images of UCP-1 in iWAT. (B) UCP-1 positive area is expressed as % of the total analyzed area in iWAT (n = 4-6). (C) Representative IHC images of UCP-1 in BAT. (D) UCP-1 positive area is expressed as % of the total analyzed area in BAT (n = 4-6). * *p* < 0.05; ** *p* < 0.01; *** *p* < 0.001 compared to HK within the same genotype. Genotype effects are shown as *p*-values above the corresponding figures (*p* < .05). Data are shown as mean ± SE. Scale bars, 200 µm.

### Tumor-induced increases in energy expenditure (EE) are not prevented by ghrelin

Tumor implantation increased EE and this difference was of greater magnitude in *Ghsr^−/−^* animals when the heat production was adjusted for lean body mass (LBM, Fig 5 A-C; endpoint EE normalized to baseline level, genotype effect: *p* = 0.013; average EE at endpoint, genotype effect: *p* = 0.010). We also analyzed the raw EE data (kcal/h) by analysis of covariance (ANCOVA) with LBM as a covariate as recommended by Tschop et al. (Tschop, Speakman et al., 2011). A significant strain difference (*p* = 0.001) was also detected using this method where *Ghsr^−/−^* mice showed higher EE levels in response to LLC tumor implantation when compared to *Ghsr^+/+^*. Animals co-administered ghrelin were not statistically different from vehicle-treated, tumor-bearing animals. Tumor implantation also decreased spontaneous locomotor activity in both genotypes and ghrelin administration did not prevent these changes (Fig 5 D-F). The respiratory quotient (RQ), was significantly decreased by tumor implantation and was not affected by genotype or ghrelin administration (Fig 5 G-I).

**Figure 5.**
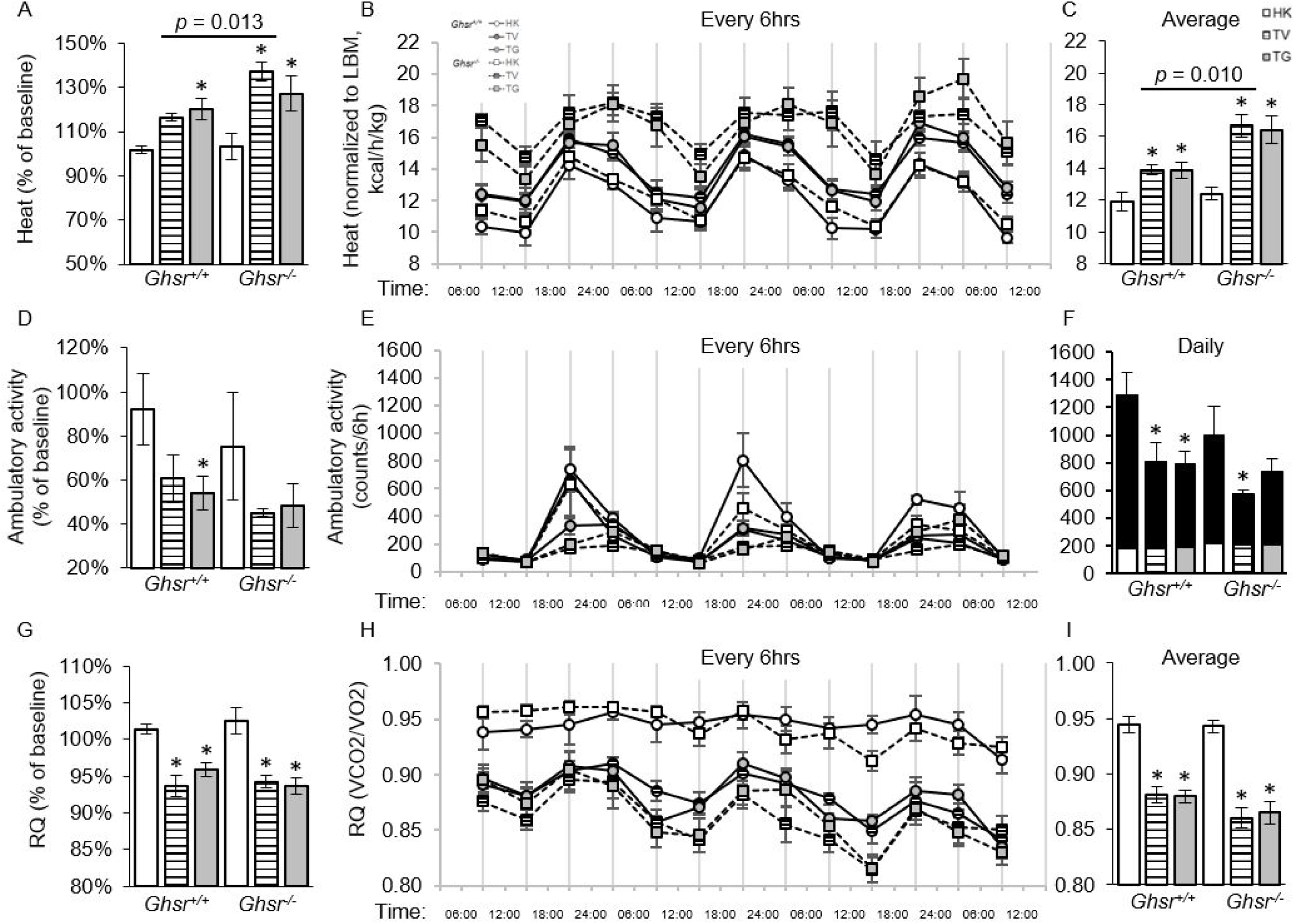
Indirect calorimetry measurements by CLAMS. HK: heat-killed + vehicle; TV: tumor + vehicle; TG: tumor + ghrelin. (A-C) Energy expenditure adjusted by LBM is expressed (A) compared to the baseline; (B) every 6 hours; and (C) average of every 6 hours. (D-F) Ambulatory activity is expressed (D) compared to baseline; (E) every 6 hours; and (F) daily (black areas represent night activity in each group). (G-I) Respiratory Quotient (RQ) is expressed (G) compared to baseline; (H) every 6 hours; and (I) average of every 6 hours. \**p*<0.05 compared to HK within the same genotype. Genotype effects are shown in *p*-values above the corresponding figures (*p* < 0.05). N = 4 for HK groups and N = 6 for the rest of the groups. Data are shown as mean ± SE.

## DISCUSSION

Adipose tissue atrophy is a central component of the cancer anorexia and cachexia syndrome (CACS) leading to increased morbidity and mortality (Das, Eder et al., 2011). Recently, emerging roles for inflammation, WAT browning and increased BAT thermogenesis have been demonstrated in this setting (Daas, Rizeq et al., 2018, Dalal, 2019, Han, Meng et al., 2018, Kir, White et al., 2014, Kliewer, Ke et al., 2015, Petruzzelli et al., 2014, Rohm, Schafer et al., 2016, Rohm, Zeigerer et al., 2019, Wang, Zhu et al., 2019); however, the pathways involved and their potential as therapeutic targets are not well-known. Ghrelin and agonists of its only known receptor, GHSR-1a, show potential to ameliorate CACS at least in part by preventing fat atrophy, but the specific mechanisms mediating these effects have not been fully characterized. Given that there are no FDA-approved treatments for cancer cachexia and that several clinical trials targeting this pathway have failed to meet their primary endpoints (Garcia et al., 2015, Temel, Abernethy et al., 2016), there is a pressing need to improve our understanding of the mechanisms of action of ghrelin in this setting. In this study we show that ghrelin prevents LLC tumor-induced weight loss, fat atrophy and WAT inflammation without affecting tumor-induced BAT inflammation, WAT browning, and increased BAT uncoupling and whole-body energy expenditure. We confirmed that its orexigenic effects are GHSR-1a-dependent, and also show that other novel GHSR-1a-independent mechanisms are involved given the partial improvements in fat atrophy and WAT inflammation seen in ghrelin-treated, *Ghsr^−/−^* animals. Also, this is the first report of macrophages as the source of IL-6 and TNF in both WAT and BAT in the setting of CACS.

Weight loss and survival rates are correlated with IL-6 levels in cancer patients (Garcia et al., 2005, Moses, Maingay et al., 2009, Scott, McMillan et al., 1996). These observations and several mechanistic studies support the premise that inflammation plays a central role in CACS. Increases in IL-1β and TNF contribute to anorexia (Baracos, Martin et al., 2018, Braun, Zhu et al., 2011, Khatib, Gaidhane et al., 2018), and TNF and IL-6 promote lipolysis and inhibit lipogenesis in WAT leading to weight loss (Fearon, Glass et al., 2012, Han et al., 2018, Jeanson, Carriere et al., 2015, Jung & Choi, 2014, Ruan, Hacohen et al., 2002). In non-cancer settings, one third of the circulating IL-6 is produced by WAT (Mohamed-Ali, Goodrick et al., 1997) and most of this WAT-derived IL-6 comes from the stroma-vascular fraction composed of endothelial cells, monocytes/macrophages, myocytes, and fibroblasts (Fain, Madan et al., 2004), although it can also be derived from adipocytes (Fain, 2006). Macrophages in WAT are known to be the source of proinflammatory cytokines in conditions leading to AT hypertrophy including obesity (Di Gregorio, Yao-Borengasser et al., 2005, Divoux, Tordjman et al., 2010, Lumeng, Deyoung et al., 2007) but this has not been previously shown in CACS. Here we show that LLC tumor implantation induces an increase in inflammatory cytokines in circulation as well as in BAT and WAT. Moreover, these AT cytokines appear to be derived exclusively from macrophages residing in these tissues. Adipose tissue atrophy in cancer patients with CACS has been associated with an increase in subcutaneous AT macrophages (Batista, Henriques et al., 2016, de Matos-Neto, Lima et al., 2015, Henriques, Sertie et al., 2017) and tissue inflammation (Batista, Olivan et al., 2013, de Matos-Neto et al., 2015, Henriques et al., 2017). Although, macrophage infiltration has also been described in WAT from tumor-bearing rodents (Henriques et al., 2017, Machado, Costa Rosa et al., 2004, Petruzzelli et al., 2014), to our knowledge this is the first report of macrophages as the source of pro-inflammatory cytokines in adipose tissue in CACS. These findings may explain why AT remains an important source of pro-inflammatory cytokines even when the adipocyte mass is significantly reduced in this setting. Also, this may be clinically relevant to cancer patients since knowing the source of inflammation may allow us to target these pathways more effectively (Henriques, Lopes et al., 2018).

Previously, we have shown that activation of GHSR-1a by ghrelin or GHSR-1a agonists (GHS) increases food intake and body weight (13, 39, 40). Our group and others also have shown that ghrelin reduces fat oxidation and lipolysis and increases lipogenesis and adiposity in a rodent model of cisplatin-induced cachexia by a combination of food intake-dependent and independent mechanisms (Chen et al., 2015, Garcia et al., 2013b, Porporato, Filigheddu et al., 2013). Ghrelin is thought to have anti-inflammatory effects in other settings (Deboer, Zhu et al., 2008, Dixit, Schaffer et al., 2004, Tsubouchi, Yanagi et al., 2014) but this is not yet clear in CACS. Some reports suggest an anti-inflammatory effect of native ghrelin administration, but this was not confirmed in other studies using GHSR-1a agonists (Chen et al., 2015, Garcia, Friend et al., 2013a). In the current study, we report that ghrelin modulates inflammation in a tissue-specific manner. Ghrelin did not prevent tumor-induced increases in circulating inflammatory cytokines or in BAT IL-1β or MCP-1 protein levels. However, it mitigated LLC-induced inflammation in WAT. This effect was seen in both genotypes although it was clearer in wild type animals partly because *Ghsr^−/−^* mice appear to be resistant to tumor-induced inflammation. GHSR-1a is not expressed in adipocytes (Sun, Garcia et al., 2007) but is present in macrophages (Ma, Lin et al., 2013) and our findings are consistent with a previous report showing that old, non-tumor-bearing *Ghsr^−/−^* mice have reduced macrophage infiltration, a shift on macrophage differentiation towards a more anti-inflammatory phenotype, and decreased inflammation in adipose tissue (Lin, Lee et al., 2016). However, a GHSR-1a-independent effect of ghrelin on macrophages is also possible as it has been proposed in other settings (Avallone, Demers et al., 2006, Bulgarelli, Tamiazzo et al., 2009, Lucchi, Costa et al., 2017). Taken together, our data is consistent with a WAT-specific, anti-inflammatory effect of ghrelin that is partly GHSR-1a dependent. This is clinically relevant as GHSR-1a agonists are in clinical development for CACS and their effect on these GHSR-1a independent pathways is not known (Garcia et al., 2015). Also, the differences we report between serum, WAT and BAT levels underscore the limitations of relying exclusively on circulating cytokine levels when trying to determine the potential role of inflammation in other tissues.

Energy expenditure is an important mechanism in the regulation of body weight and is increased in CACS (Garcia et al., 2013a, Kir, Komaba et al., 2016, Rohm et al., 2019). Factors contributing to EE include physical activity and resting EE (REE) (Silver, Dietrich et al., 2007, Vazeille, Jouinot et al., 2017) and adipose tissue can lead to an increase in REE by uncoupling oxidative phosphorylation in mitochondria thereby releasing heat through activation of a proton leak (Nicholls, 1976, Okamatsu-Ogura, Kitao et al., 2007). In WAT, browning has been noted in multiple cancer cachexia models with adipocytes showing an upregulation of the main regulator of thermogenesis, UCP1 (Dong, Lin et al., 2018, Vaitkus & Celi, 2017). In BAT, increased thermogenesis has been reported in cachectic animals (Kir et al., 2014) independently of decreased food intake or their ability to maintain their body temperature (Tsoli, Moore et al., 2012). Proinflammatory cytokines have been suggested as key drivers of WAT browning (Han et al., 2018, Petruzzelli et al., 2014) and of BAT thermogenesis through activation of sympathetic nervous system or targeting BAT directly (Arruda, Milanski et al., 2010, Dascombe, Rothwell et al., 1989, Li, Klein et al., 2002, Tsoli et al., 2012). Here we show that LLC-tumor implantation led to an increase in total EE in spite of a significant decrease in physical activity, suggesting an increase in REE. This was associated with an increase in UCP-1 expression in WAT (browning) and in BAT. Moreover, these effects were more marked in *Ghsr^−/−^* mice suggesting a protective role of GHSR-1a in this setting. These results agree with previous reports in aged, non-tumor-bearing *Ghsr^−/−^* showing higher levels of thermogenesis and energy expenditure when compared to aged-matched, wild-type mice (Lin, Saha et al., 2011). The effect of ghrelin or GHSR1a agonists on energy expenditure is unclear with some studies showing a decrease in EE (Borner, Loi et al., 2016, Villars, Pietra et al., 2017) while others showed no effect (Adachi, Takiguchi et al., 2010, Tschop, Smiley et al., 2000, Vestergaard, Djurhuus et al., 2008). In this study, we did not see a significant effect of ghrelin on preventing LLC-induced fat browning, BAT thermogenesis, increased REE or decreased physical activity in the setting of CACS despite the fact that ghrelin prevented fat and weight loss and anorexia. We hypothesize that differences in the models, route of administration and treatment regimen and agents used (LLC mice vs. C26 mice or hepatoma model in rats, administration via s.q. vs. oral gavage vs. osmotic mini pump, ghrelin vs. GHSR1a agonists) could account for these discrepancies. More studies will be needed to test this hypothesis.

Macrophage infiltration contributes to the high levels of inflammatory cytokines (TNF, IL-6, and IL-1β) in BAT in conditions associated with AT hypertrophy such as high fat diet (Roberts-Toler, O’Neill et al., 2015, van den Berg, van Dam et al., 2017) or obesity (Alcala, Calderon-Dominguez et al., 2017, Calderon-Dominguez, Mir et al., 2016). In CACS the aforementioned tumor-induced inflammation is thought to play an important role in BAT thermogenesis (Petruzzelli et al., 2014, Tsoli et al., 2012); however, the source of inflammation in BAT is not known. Similar to WAT, we found that BAT IL-6 and TNF come exclusively from macrophages in the setting of cachexia. However, their expression in BAT were lower than in WAT and no significant changes were found in response to tumor implantation or ghrelin. We found a significant tumor-effect on increasing IL-1β levels in BAT although ghrelin did not prevent this increase, suggesting tissue-specific differences in inflammation between BAT and WAT in response to tumor and ghrelin. Taken together, these results are important because they show that tumor-induced WAT browning and BAT thermogenesis are associated with significant increases in REE and appear to be independent of inflammation given that downregulating inflammation does not prevent uncoupling in WAT and that BAT IL6 and TNF levels were not upregulated upon tumor implantation. In addition, our data suggests that WAT is a significant source of inflammatory cytokines, which express the highest levels of IL-1β, IL-6, and TNF when compared to BAT and circulating levels.

There were limitations to our approach. This study was not set up to establish the safety of ghrelin administration in the setting of cancer. Nevertheless, none of the studies published to date using ghrelin or GHSR-1a agonists in mice or humans have shown an increase in tumor progression (Sever, White et al., 2016). Also, the experiments were not designed to characterize other mechanisms contributing to the protective role of GHSR-1a in this setting. Lastly, our data suggest that there is an alternative receptor for ghrelin although identification of this receptor remains elusive and is the focus of other studies.

In summary, ghrelin prevents LLC tumor-induced body weight and fat loss by a combination of GHSR-1a-dependent mechanisms including preventing anorexia, and other mechanisms that are partly GHSR-1a-independent. The increase in inflammation in AT induced by tumor implantation is prevented by ghrelin only in WAT; however, tumor-induced WAT browning, and increased BAT inflammation, uncoupling and whole body energy expenditure are not prevented by ghrelin even when the presence of GHSR-1a appears to contribute to maintaining energy balance in this setting. Tumor-induced WAT browning and BAT thermogenesis are associated with significant increases in REE and these seem to be independent of inflammation given that downregulating it does not prevent these changes. These results are clinically relevant because they show that ghrelin ameliorates WAT inflammation, fat atrophy and anorexia in CACS in spite of not having a discernible effect on energy expenditure, WAT browning or BAT inflammation and thermogenesis. Our data fills an important gap in the knowledge regarding the mechanisms of action of ghrelin in the setting of cancer cachexia and should inform the design of future preclinical and clinical studies targeting this pathway.

## METHODS

### Animals

Five to seven-month-old male C57BL/6J growth hormone (GH) secretagogue receptor wild type (*Ghsr^+/+^*) and knockout (*Ghsr^−/−^*) congenic mice were used for all experiments. Briefly the *Ghsr^+/+^* and *Ghsr^−/−^* mice were originally from Dr. Roy G. Smith Ph.D’s laboratory (Sun, Butte et al., 2008) and the *Ghsr^−/−^* mice were backcrossed with C57BL/6J for at least 10 generations to minimize selective genetic traits. The mice used in the study were off springs of these congenic mice and were bred in the Animal Research Facilities in Veterans Affairs Puget Sound Health Care System. Mice were individually housed, acclimated to their cages and human handling for 1 week before the experiments and maintained on a 12/12 light/dark cycle (lights on at 6AM). All experiments were conducted with the approval of the Institutional Animal Care and Use Committee at VA Puget Sound Health Care System and were in compliance with the NIH Guidelines for Use and Care of Laboratory Animals. Sample sizes of each experiment are shown in the figure legends.

### Tumor implantation and ghrelin administration

The procedures of tumor implantation (TI) and ghrelin intervention were described previously (Chen et al., 2015). In brief, mice were injected subcutaneously (s.q.) with Lewis lung carcinoma (LLC) cells (1 × 10^6^ cells, CRL1642, American Type Culture Collection, Manassas, VA) into the right flank or with equal volume and number of heat-killed LLC cells (HK). Approximately 7 days after tumor implantation (TI), when the tumor was palpable (∼1cm in diameter), the tumor-bearing mice were treated with either acylated ghrelin (AS-24160, Anaspect, Fremont, CA) at a dose of 0.8 mg/kg or vehicle (0.9% sodium chloride, 8881570121, COVIDIEN, Dublin, Ireland), s.q., twice daily, while mice in HK group received vehicle (saline, same volume), s.q., twice daily for two weeks. Mice were euthanized by CO_2_ on Day 21 after TI, approximately 2 weeks after TN. Blood samples were collected and then processed into plasma. Fat pads including iWAT, eWAT, and BAT, as well as tumors were collected during dissection. The timeline of the study is demonstrated in Supplemental Fig. 5.

### Body weight, food intake, and body composition

Body weight and food intake were assessed daily starting before TI (baseline) until endpoint. Parameters of body composition, including LBM and fat mass (FM) were measured by nuclear magnetic resonance (NMR, Bruker optics, The Woodlands, TX) and identified at the baseline before tumor implantation, when tumor was noted, and 2 weeks after tumor noted before terminating the experiment (endpoint).

### Comprehensive laboratory animal monitoring system (CLAMS™)

The Comprehensive Laboratory Animal Monitoring System (CLAMS™, Columbus Instruments, Columbus, OH) was used to identify metabolic parameters of the animals as we previously described (Guillory, Chen et al., 2017). *Ghsr^+/+^* and *Ghsr^−/−^* mice were individually housed in CLAMS cages for 96 hours before TI as well as at the endpoint (see the Supplemental Fig. 5, timeline for the study). The first 12 hours of CLAMS was considered as the acclimation phase and the data for the next 72 hours were analyzed. Oxygen consumption (VO_2_) (mL/h), carbon dioxide production (VCO_2_) (mL/h), and locomotor activity (infrared beam-break counts) were recorded automatically by the CLAMS system every 20 min. The respiratory exchange ratio (RQ) and energy expenditure (EE, or heat generation) were calculated from VO_2_ and VCO_2_ gas exchange data as follows: RQ = VCO_2_/VO_2_ and EE = (3.815 + 1.232 × RQ) × VO_2_, respectively. Energy expenditure was then normalized to LBM for statistical analysis using two-way analysis of variance (ANOVA). Alternatively, we also analyzed EE value by ANCOVA with LBM as a covariate. Locomotor activity was measured on x- and z-axes by the counts of beam-breaks during the recording period. The data shown in the results was summarized as the mean of every 6 hours in a 72-hour-period.

### Electrochemiluminescence immunoassay

Inflammatory cytokines IL-1β, IL-6, and TNF-α and macrophage marker MCP-1 in iWAT, BAT, and serum were detected by U-PLEX Biomarker Group1 (ms) Assays which are developed by Meso Scale Diagnostics (K15069L-1, MSD, Rockville, MD). A protocol provided by manufacturer was used for this assay. In brief, each plate was prepared by overnight coating with the multiplex coating solution at 4 °C, which contained linker-coupled biotinylated antibodies. Standards and serum samples were diluted with Diluent 41 into 2-fold and loaded onto the coated plate on the next day. For iWAT and BAT samples, 150ug of the protein lysate was diluted with Diluent 41 and loaded onto each well. The plate was incubated at room temperature (RT) with shaking for 2h followed by 3 times of wash in phosphate buffered saline with .05% Tween 20 (PBS/T). Sulfo-tag labeled detection antibody was then added to plates and incubated for 2.5h. After another 3 washes in PBS/T, Read Buffer T(2x) was added and the plate was read on MSD Sector Imager (MSD).

### Immunohistochemistry

The iWAT and BAT were mounted with OCT (VWR 25608-930, VWR, Radnor, PA) and flash frozen in liquid nitrogen-chilled isopentane immediately after tissue collection. The OCT-mounted iWAT and BAT blocks were sliced at 14μm using a Cryostat (Leica CM3050S, Nussloch, Germany) at −40°C. Before the process of staining, slides were dehydrated at RT for 30 minutes followed by incubating in methanol for 15 minutes at −20 °C. To identify the colocalization of F4/80 and IL-6 or TNFα in iWAT and BAT, slides were blocked with 10% donkey serum for 1 hour at RT and followed by incubating in primary antibodies (F4/80 Monoclonal Antibody 1:100, MF48000, Thermo Fisher Scientific; Anti-IL-6 antibody 1:100, ab6672, Abcam; TNF alpha monoclonal antibody, FITC, eBioscience™ 1:200, 11-7349-82, Thermo Fisher Scientific) at 4°C for overnight. After 3 washes in PBS, the slides were incubated by the corresponding secondary antibodies (Alexa Fluor 594 donkey anti-rat IgG, A21209, or Alexa Fluor 488 donkey anti-rat IgG, A21208, for F4/80; Texas Red goat anti-rabbit IgG, T-2767, for IL-6) for 2 hours at RT and followed by incubating in 1:1000 DAPI (62248, Thermo Fisher Scientific) in PBS for 1min. The slides were then mounted by Prolong Gold AntiFade reagent (P36930, Thermo Fisher Scientific) with coverslips. To identify UCP1 in iWAT and BAT, slides were incubated with 3% hydrogen peroxide (323381, Sigma-Aldrich, St. Louis, MO) for 30 min and then in 2.5% normal horse serum for 1hr. Then the slides were incubated with UCP1 Polyclonal Antibody (PA1-24894, Thermo Fisher Scientific) diluted 1:200 in 2.5% normal horse serum at 4°C for overnight. On the following day, signals were visualized using SignalStain® Boost IHC Detection Reagent (8114, Cell Signaling) and the SignalStain® DAB Substrate kit (8059, Cell Signaling). The stained slides were dehydrated by 70%, 90%, 100% ethanol, and 100% xylene sequentially and mounted with coverslips by using Permount (SP15-100, Thermo Fisher Scientific). All stained slides were imaged by Nikon NiE microscope at 20x (iWAT) or 40x (BAT). The positive cells (immunofluorescence) or positive area (DAB stain) in the section were quantified and normalized to the total area of the section (mm^2^) using ImageJ analysis software (National Institutes of Health, http://rsb.info.nih.gov/ij/).

### Statistics

Two-way ANOVA was performed to identify differences between genotypes (*Ghsr^+/+^* vs. *Ghsr^−/−^*) across treatments (HK, TV, and TG) followed by Fisher’s LSD post hoc test. For inflammatory cytokines, Kruskal-Wallis test was performed to identify the differences between groups. For energy expenditure, ANCOVA was also used for analysis in addition to ANOVA with LBM as a covariate to identify differences between genotypes across treatments followed by Fisher’s LSD post hoc test. Values are presented in mean ± SEM. All statistical testing was performed using IBM SPSS version 18 software. Significant difference was set at *: *p* < 0.05; **: *p* < 0.01; ***: *p* < 0.001.

## ACKNOWEDGEMENTS

We would like to thank Dr. Tammy Wolden-Hanson for helping with the measures from NMR and CLAMS at the Rodent Metabolic and Behavioral Phenotyping Core at the VA Puget Sound Health Care System. We would also like to thank Dr. Rebecca Hull, Nishi Ivanov, and Daryl Hackney for providing guidance on immunohistochemistry and imaging at the Cellular and Molecular Imaging Core at the Diabetes Research Center in University of Washington. We would like to acknowledge the National Institutes of Health (NIH) National Institute of Diabetes and Digestive and Kidney Diseases funded Nutrition Obesity Research Center (DK035816) and Diabetes Research Center (P30 DK017047) at the University of Washington.

## AUTHOR CONTRIBUTIONS

HL, JL, BG, and JMG designed the study. HL, JL, PZ, JAC, JKY, YH and BA conducted experiments and acquired data. HL, JL, BG, JAC, PZ, and IL handled the mice in the study. HL, JL, BG, JAC, PZ, IL, BA, MS, and AT collected tissue. HL, JL, BA, MS, and AT analyzed data. HL, JL, and JMG wrote the manuscript. All authors reviewed and approved the final version of the manuscript.

---

Supplemental Fig. 1. Gene expression of *Ghsr* in brain, iWAT, and BAT in *Ghsr* ^+/+^ and ^−/−^ mice. Data is expressed as box-and-whisker plot showing the median (middle line), mean (middle cross), upper and lower quartiles (box), maximum and minimum (whiskers). Relative gene expression was determined by normalization to *Gapdh*. N = 4/group. *Ghsr* was only detected in brain in *Ghsr* ^+/+^ mice. No *Ghsr* expression was detected in any tissue in *Ghsr* ^−/−^ or adipose tissue in *Ghsr* ^+/+^ mice.

Supplemental Fig. 2. High resolution images of immunohistochemistry staining in iWAT. (A) Representative images of colocalization of inflammatory marker IL-6 and macrophage marker F4/80 in iWAT (IL-6 in Texas red; F4/80 in FITC green; nuclei in DAPI blue). (B) Representative images of colocalization of inflammatory marker TNF and macrophage marker F4/80 in iWAT (TNF in FITC green; F4/80 in Texas red; nuclei in DAPI blue). Positively stained inflammatory markers and colocalizations with macrophages are indicated by the white arrows. Scale bars, 100 µm.

Supplemental Fig. 3. High resolution images of immunohistochemistry staining in BAT. (A) Representative images of colocalization of inflammatory marker IL-6 and macrophage marker F4/80 in BAT (IL-6 in Texas red; F4/80 in FITC green; nuclei in DAPI blue). (B) Representative images of colocalization of inflammatory marker TNF and macrophage marker F4/80 in BAT (TNF in FITC green; F4/80 in Texas red; nuclei in DAPI blue). Positively stained inflammatory markers and colocalizations with macrophages are indicated by the white arrows. Scale bars, 100 µm.

Supplemental Fig. 4. Effects of ghrelin on LLC-induced protein-level changes in inflammation (IL-1β, IL-6, and TNF) and macrophages (MCP-1) in plasma (pg/mg, n = 11-14). *, **: different than HK within the same genotype (*: p < .05; **: p < .01). Genotype effects are shown in p-values above the corresponding figures (p < .05). Data are shown as mean ± SE.

Supplemental Fig. 4. Timeline of current study. *Ghsr^+/+^* and *^−/−^* mice were injected with LLC (T, 1 × 106 cells, s.q.) into the right flank or with equal volume and number of heat-killed LLC cells (HK). Approximately 7 days after tumor implantation, when the tumor was palpable (day 0), the tumor-bearing mice were treated with either acylated ghrelin, 0.8 mg/kg (TG) or vehicle (0.9% sodium chloride, TV), s.q., twice daily, while mice in HK group received vehicle (saline, same volume), s.q., twice daily for two weeks. Body composition were identified by NMR before tumor implantation (7 days before tumor noted, baseline) and weekly till the endpoint. All the mice were individually housed in CLAMS cages for 96 hours before TI (11-7 days before tumor noted, baseline) as well as at the endpoint (day 10-14 after tumor noted).

## REFERENCES

Adachi S, Takiguchi S, Okada K, Yamamoto K, Yamasaki M, Miyata H, Nakajima K, Fujiwara Y, Hosoda H, Kangawa K, Mori M, Doki Y (2010) Effects of ghrelin administration after total gastrectomy: a prospective, randomized, placebo-controlled phase II study. Gastroenterology 138: 1312–20

Alcala M, Calderon-Dominguez M, Bustos E, Ramos P, Casals N, Serra D, Viana M, Herrero L (2017) Increased inflammation, oxidative stress and mitochondrial respiration in brown adipose tissue from obese mice. Sci Rep 7: 16082

Arruda AP, Milanski M, Romanatto T, Solon C, Coope A, Alberici LC, Festuccia WT, Hirabara SM, Ropelle E, Curi R, Carvalheira JB, Vercesi AE, Velloso LA (2010) Hypothalamic actions of tumor necrosis factor alpha provide the thermogenic core for the wastage syndrome in cachexia. Endocrinology 151: 683–94

Avallone R, Demers A, Rodrigue-Way A, Bujold K, Harb D, Anghel S, Wahli W, Marleau S, Ong H, Tremblay A (2006) A growth hormone-releasing peptide that binds scavenger receptor CD36 and ghrelin receptor up-regulates sterol transporters and cholesterol efflux in macrophages through a peroxisome proliferator-activated receptor gamma-dependent pathway. Mol Endocrinol 20: 3165–78

Baracos VE, Martin L, Korc M, Guttridge DC, Fearon KCH (2018) Cancer-associated cachexia. Nat Rev Dis Primers 4: 17105

Batista ML, Jr., Henriques FS, Neves RX, Olivan MR, Matos-Neto EM, Alcantara PS, Maximiano LF, Otoch JP, Alves MJ, Seelaender M (2016) Cachexia-associated adipose tissue morphological rearrangement in gastrointestinal cancer patients. J Cachexia Sarcopenia Muscle 7: 37–47

Batista ML, Jr., Olivan M, Alcantara PS, Sandoval R, Peres SB, Neves RX, Silverio R, Maximiano LF, Otoch JP, Seelaender M (2013) Adipose tissue-derived factors as potential biomarkers in cachectic cancer patients. Cytokine 61: 532–9

Borner T, Loi L, Pietra C, Giuliano C, Lutz TA, Riediger T (2016) The ghrelin receptor agonist HM01 mimics the neuronal effects of ghrelin in the arcuate nucleus and attenuates anorexia-cachexia syndrome in tumor-bearing rats. Am J Physiol Regul Integr Comp Physiol 311: R89–96

Braun TP, Zhu X, Szumowski M, Scott GD, Grossberg AJ, Levasseur PR, Graham K, Khan S, Damaraju S, Colmers WF, Baracos VE, Marks DL (2011) Central nervous system inflammation induces muscle atrophy via activation of the hypothalamic-pituitary-adrenal axis. J Exp Med 208: 2449–63

Bulgarelli I, Tamiazzo L, Bresciani E, Rapetti D, Caporali S, Lattuada D, Locatelli V, Torsello A (2009) Desacyl-ghrelin and synthetic GH-secretagogues modulate the production of inflammatory cytokines in mouse microglia cells stimulated by beta-amyloid fibrils. J Neurosci Res 87: 2718–27

Calderon-Dominguez M, Mir JF, Fucho R, Weber M, Serra D, Herrero L (2016) Fatty acid metabolism and the basis of brown adipose tissue function. Adipocyte 5: 98–118

Chen JA, Splenser A, Guillory B, Luo J, Mendiratta M, Belinova B, Halder T, Zhang G, Li YP, Garcia JM (2015) Ghrelin prevents tumour- and cisplatin-induced muscle wasting: characterization of multiple mechanisms involved. J Cachexia Sarcopenia Muscle 6: 132–43

Daas SI, Rizeq BR, Nasrallah GK (2018) Adipose tissue dysfunction in cancer cachexia. J Cell Physiol 234: 13–22

Dalal S (2019) Lipid metabolism in cancer cachexia. Ann Palliat Med 8: 13–23

Das SK, Eder S, Schauer S, Diwoky C, Temmel H, Guertl B, Gorkiewicz G, Tamilarasan KP, Kumari P, Trauner M, Zimmermann R, Vesely P, Haemmerle G, Zechner R, Hoefler G (2011) Adipose triglyceride lipase contributes to cancer-associated cachexia. Science 333: 233–8

Dascombe MJ, Rothwell NJ, Sagay BO, Stock MJ (1989) Pyrogenic and thermogenic effects of interleukin 1 beta in the rat. Am J Physiol 256: E7–11

de Matos-Neto EM, Lima JD, de Pereira WO, Figueredo RG, Riccardi DM, Radloff K, das Neves RX, Camargo RG, Maximiano LF, Tokeshi F, Otoch JP, Goldszmid R, Camara NO, Trinchieri G, de Alcantara PS, Seelaender M (2015) Systemic Inflammation in Cachexia - Is Tumor Cytokine Expression Profile the Culprit? Front Immunol 6: 629

Deboer MD, Zhu X, Levasseur PR, Inui A, Hu Z, Han G, Mitch WE, Taylor JE, Halem HA, Dong JZ, Datta R, Culler MD, Marks DL (2008) Ghrelin treatment of chronic kidney disease: improvements in lean body mass and cytokine profile. Endocrinology 149: 827–35

Deshmane SL, Kremlev S, Amini S, Sawaya BE (2009) Monocyte chemoattractant protein-1 (MCP-1): an overview. J Interferon Cytokine Res 29: 313–26

Dewys WD, Begg C, Lavin PT, Band PR, Bennett JM, Bertino JR, Cohen MH, Douglass HO, Jr., Engstrom PF, Ezdinli EZ, Horton J, Johnson GJ, Moertel CG, Oken MM, Perlia C, Rosenbaum C, Silverstein MN, Skeel RT, Sponzo RW, Tormey DC (1980) Prognostic effect of weight loss prior to chemotherapy in cancer patients. Eastern Cooperative Oncology Group. Am J Med 69: 491–7

Di Gregorio GB, Yao-Borengasser A, Rasouli N, Varma V, Lu T, Miles LM, Ranganathan G, Peterson CA, McGehee RE, Kern PA (2005) Expression of CD68 and macrophage chemoattractant protein-1 genes in human adipose and muscle tissues: association with cytokine expression, insulin resistance, and reduction by pioglitazone. Diabetes 54: 2305–13

Divoux A, Tordjman J, Lacasa D, Veyrie N, Hugol D, Aissat A, Basdevant A, Guerre-Millo M, Poitou C, Zucker JD, Bedossa P, Clement K (2010) Fibrosis in human adipose tissue: composition, distribution, and link with lipid metabolism and fat mass loss. Diabetes 59: 2817–25

Dixit VD, Schaffer EM, Pyle RS, Collins GD, Sakthivel SK, Palaniappan R, Lillard JW, Jr., Taub DD (2004) Ghrelin inhibits leptin- and activation-induced proinflammatory cytokine expression by human monocytes and T cells. J Clin Invest 114: 57–66

Dong M, Lin J, Lim W, Jin W, Lee HJ (2018) Role of brown adipose tissue in metabolic syndrome, aging, and cancer cachexia. Front Med 12: 130–138

Fain JN (2006) Release of interleukins and other inflammatory cytokines by human adipose tissue is enhanced in obesity and primarily due to the nonfat cells. Vitam Horm 74: 443–477

Fain JN, Madan AK, Hiler ML, Cheema P, Bahouth SW (2004) Comparison of the release of adipokines by adipose tissue, adipose tissue matrix, and adipocytes from visceral and subcutaneous abdominal adipose tissues of obese humans. Endocrinology 145: 2273–82

Fearon K, Strasser F, Anker SD, Bosaeus I, Bruera E, Fainsinger RL, Jatoi A, Loprinzi C, MacDonald N, Mantovani G, Davis M, Muscaritoli M, Ottery F, Radbruch L, Ravasco P, Walsh D, Wilcock A, Kaasa S, Baracos VE (2011) Definition and classification of cancer cachexia: an international consensus. Lancet Oncol 12: 489–95

Fearon KC, Glass DJ, Guttridge DC (2012) Cancer cachexia: mediators, signaling, and metabolic pathways. Cell Metab 16: 153–66

Fouladiun M, Korner U, Bosaeus I, Daneryd P, Hyltander A, Lundholm KG (2005) Body composition and time course changes in regional distribution of fat and lean tissue in unselected cancer patients on palliative care--correlations with food intake, metabolism, exercise capacity, and hormones. Cancer 103: 2189–98

Garcia JM, Boccia RV, Graham CD, Yan Y, Duus EM, Allen S, Friend J (2015) Anamorelin for patients with cancer cachexia: an integrated analysis of two phase 2, randomised, placebo-controlled, double-blind trials. Lancet Oncol 16: 108–16

Garcia JM, Friend J, Allen S (2013a) Therapeutic potential of anamorelin, a novel, oral ghrelin mimetic, in patients with cancer-related cachexia: a multicenter, randomized, double-blind, crossover, pilot study. Support Care Cancer 21: 129–37

Garcia JM, Garcia-Touza M, Hijazi RA, Taffet G, Epner D, Mann D, Smith RG, Cunningham GR, Marcelli M (2005) Active ghrelin levels and active to total ghrelin ratio in cancer-induced cachexia. J Clin Endocrinol Metab 90: 2920–6

Garcia JM, Scherer T, Chen JA, Guillory B, Nassif A, Papusha V, Smiechowska J, Asnicar M, Buettner C, Smith RG (2013b) Inhibition of cisplatin-induced lipid catabolism and weight loss by ghrelin in male mice. Endocrinology 154: 3118–29

Guillory B, Chen JA, Patel S, Luo JH, Splenser A, Mody A, Ding M, Baghaie S, Anderson B, Lankova B, Halder T, Hernandez Y, Garcia JM (2017) Deletion of ghrelin prevents aging-associated obesity and muscle dysfunction without affecting longevity. Aging Cell 16: 859–869

Han J, Meng Q, Shen L, Wu G (2018) Interleukin-6 induces fat loss in cancer cachexia by promoting white adipose tissue lipolysis and browning. Lipids Health Dis 17: 14

Henriques F, Lopes MA, Franco FO, Knobl P, Santos KB, Bueno LL, Correa VA, Bedard AH, Guilherme A, Birbrair A, Peres SB, Farmer SR, Batista ML, Jr. (2018) Toll-Like Receptor-4 Disruption Suppresses Adipose Tissue Remodeling and Increases Survival in Cancer Cachexia Syndrome. Sci Rep 8: 18024

Henriques FS, Sertie RAL, Franco FO, Knobl P, Neves RX, Andreotti S, Lima FB, Guilherme A, Seelaender M, Batista ML, Jr. (2017) Early suppression of adipocyte lipid turnover induces immunometabolic modulation in cancer cachexia syndrome. FASEB J 31: 1976–1986

Jeanson Y, Carriere A, Casteilla L (2015) A New Role for Browning as a Redox and Stress Adaptive Mechanism? Front Endocrinol (Lausanne) 6: 158

Jung UJ, Choi MS (2014) Obesity and its metabolic complications: the role of adipokines and the relationship between obesity, inflammation, insulin resistance, dyslipidemia and nonalcoholic fatty liver disease. Int J Mol Sci 15: 6184–223

Khatib MN, Gaidhane A, Gaidhane S, Quazi ZS (2018) Ghrelin as a Promising Therapeutic Option for Cancer Cachexia. Cell Physiol Biochem 48: 2172–2188

Kir S, Komaba H, Garcia AP, Economopoulos KP, Liu W, Lanske B, Hodin RA, Spiegelman BM (2016) PTH/PTHrP Receptor Mediates Cachexia in Models of Kidney Failure and Cancer. Cell Metab 23: 315–23

Kir S, White JP, Kleiner S, Kazak L, Cohen P, Baracos VE, Spiegelman BM (2014) Tumour-derived PTH-related protein triggers adipose tissue browning and cancer cachexia. Nature 513: 100–4

Kliewer KL, Ke JY, Tian M, Cole RM, Andridge RR, Belury MA (2015) Adipose tissue lipolysis and energy metabolism in early cancer cachexia in mice. Cancer Biol Ther 16: 886–97

Kojima M, Hosoda H, Date Y, Nakazato M, Matsuo H, Kangawa K (1999) Ghrelin is a growth-hormone-releasing acylated peptide from stomach. Nature 402: 656–60

Kos K, Harte AL, O’Hare PJ, Kumar S, McTernan PG (2009) Ghrelin and the differential regulation of des-acyl (DSG) and oct-anoyl ghrelin (OTG) in human adipose tissue (AT). Clin Endocrinol (Oxf) 70: 383–9

Lerner L, Hayes TG, Tao N, Krieger B, Feng B, Wu Z, Nicoletti R, Chiu MI, Gyuris J, Garcia JM (2015) Plasma growth differentiation factor 15 is associated with weight loss and mortality in cancer patients. J Cachexia Sarcopenia Muscle 6: 317–24

Li G, Klein RL, Matheny M, King MA, Meyer EM, Scarpace PJ (2002) Induction of uncoupling protein 1 by central interleukin-6 gene delivery is dependent on sympathetic innervation of brown adipose tissue and underlies one mechanism of body weight reduction in rats. Neuroscience 115: 879–889

Lin L, Lee JH, Buras ED, Yu K, Wang R, Smith CW, Wu H, Sheikh-Hamad D, Sun Y (2016) Ghrelin receptor regulates adipose tissue inflammation in aging. Aging (Albany NY) 8: 178–91

Lin L, Saha PK, Ma X, Henshaw IO, Shao L, Chang BH, Buras ED, Tong Q, Chan L, McGuinness OP, Sun Y (2011) Ablation of ghrelin receptor reduces adiposity and improves insulin sensitivity during aging by regulating fat metabolism in white and brown adipose tissues. Aging Cell 10: 996–1010

Lucchi C, Costa AM, Giordano C, Curia G, Piat M, Leo G, Vinet J, Brunel L, Fehrentz JA, Martinez J, Torsello A, Biagini G (2017) Involvement of PPARgamma in the Anticonvulsant Activity of EP-80317, a Ghrelin Receptor Antagonist. Front Pharmacol 8: 676

Lumeng CN, Deyoung SM, Bodzin JL, Saltiel AR (2007) Increased inflammatory properties of adipose tissue macrophages recruited during diet-induced obesity. Diabetes 56: 16–23

Ma X, Lin L, Yue J, Pradhan G, Qin G, Minze LJ, Wu H, Sheikh-Hamad D, Smith CW, Sun Y (2013) Ghrelin receptor regulates HFCS-induced adipose inflammation and insulin resistance. Nutr Diabetes 3: e99

Machado AP, Costa Rosa LF, Seelaender MC (2004) Adipose tissue in Walker 256 tumour-induced cachexia: possible association between decreased leptin concentration and mononuclear cell infiltration. Cell Tissue Res 318: 503–14

Michaud A, Boulet MM, Veilleux A, Noel S, Paris G, Tchernof A (2014) Abdominal subcutaneous and omental adipocyte morphology and its relation to gene expression, lipolysis and adipocytokine levels in women. Metabolism 63: 372–81

Mohamed-Ali V, Goodrick S, Rawesh A, Katz DR, Miles JM, Yudkin JS, Klein S, Coppack SW (1997) Subcutaneous adipose tissue releases interleukin-6, but not tumor necrosis factor-alpha, in vivo. J Clin Endocrinol Metab 82: 4196–200

Moses AG, Maingay J, Sangster K, Fearon KC, Ross JA (2009) Pro-inflammatory cytokine release by peripheral blood mononuclear cells from patients with advanced pancreatic cancer: relationship to acute phase response and survival. Oncol Rep 21: 1091–5

Muller TD, Tschop MH (2013) Ghrelin - a key pleiotropic hormone-regulating systemic energy metabolism. Endocr Dev 25: 91–100

Nicholls DG (1976) The bioenergetics of brown adipose tissue mitochondria. FEBS Letters 61: 103–110

Okamatsu-Ogura Y, Kitao N, Kimura K, Saito M (2007) Brown fat UCP1 is not involved in the febrile and thermogenic responses to IL-1beta in mice. Am J Physiol Endocrinol Metab 292: E1135–9

Perez-Tilve D, Heppner K, Kirchner H, Lockie SH, Woods SC, Smiley DL, Tschop M, Pfluger P (2011) Ghrelin-induced adiposity is independent of orexigenic effects. FASEB J 25: 2814–22

Petruzzelli M, Schweiger M, Schreiber R, Campos-Olivas R, Tsoli M, Allen J, Swarbrick M, Rose-John S, Rincon M, Robertson G, Zechner R, Wagner EF (2014) A switch from white to brown fat increases energy expenditure in cancer-associated cachexia. Cell Metab 20: 433–47

Porporato PE, Filigheddu N, Reano S, Ferrara M, Angelino E, Gnocchi VF, Prodam F, Ronchi G, Fagoonee S, Fornaro M, Chianale F, Baldanzi G, Surico N, Sinigaglia F, Perroteau I, Smith RG, Sun Y, Geuna S, Graziani A (2013) Acylated and unacylated ghrelin impair skeletal muscle atrophy in mice. J Clin Invest 123: 611–22

Puigserver P, Wu Z, Park CW, Graves R, Wright M, Spiegelman BM (1998) A Cold-Inducible Coactivator of Nuclear Receptors Linked to Adaptive Thermogenesis. Cell 92: 829–839

Roberts-Toler C, O’Neill BT, Cypess AM (2015) Diet-induced obesity causes insulin resistance in mouse brown adipose tissue. Obesity (Silver Spring) 23: 1765–70

Rohm M, Schafer M, Laurent V, Ustunel BE, Niopek K, Algire C, Hautzinger O, Sijmonsma TP, Zota A, Medrikova D, Pellegata NS, Ryden M, Kulyte A, Dahlman I, Arner P, Petrovic N, Cannon B, Amri EZ, Kemp BE, Steinberg GR et al. (2016) An AMP-activated protein kinase-stabilizing peptide ameliorates adipose tissue wasting in cancer cachexia in mice. Nat Med 22: 1120–1130

Rohm M, Zeigerer A, Machado J, Herzig S (2019) Energy metabolism in cachexia. EMBO Rep 20

Rosen ED, Spiegelman BM (2014) What we talk about when we talk about fat. Cell 156: 20–44

Ruan H, Hacohen N, Golub TR, Van Parijs L, Lodish HF (2002) Tumor necrosis factor-alpha suppresses adipocyte-specific genes and activates expression of preadipocyte genes in 3T3-L1 adipocytes: nuclear factor-kappaB activation by TNF-alpha is obligatory. Diabetes 51: 1319–36

Scott HR, McMillan DC, Crilly A, McArdle CS, Milroy R (1996) The relationship between weight loss and interleukin 6 in non-small-cell lung cancer. Brit J Cancer 73: 1560–1562

Sever S, White DL, Garcia JM (2016) Is there an effect of ghrelin/ghrelin analogs on cancer? A systematic review. Endocr Relat Cancer 23: R393–409

Silver HJ, Dietrich MS, Murphy BA (2007) Changes in body mass, energy balance, physical function, and inflammatory state in patients with locally advanced head and neck cancer treated with concurrent chemoradiation after low-dose induction chemotherapy. Head Neck 29: 893–900

Smith RG, Van der Ploeg LH, Howard AD, Feighner SD, Cheng K, Hickey GJ, Wyvratt MJ, Jr., Fisher MH, Nargund RP, Patchett AA (1997) Peptidomimetic regulation of growth hormone secretion. Endocr Rev 18: 621–645

Sun Y, Butte NF, Garcia JM, Smith RG (2008) Characterization of adult ghrelin and ghrelin receptor knockout mice under positive and negative energy balance. Endocrinology 149: 843–50

Sun Y, Garcia JM, Smith RG (2007) Ghrelin and growth hormone secretagogue receptor expression in mice during aging. Endocrinology 148: 1323–9

Temel JS, Abernethy AP, Currow DC, Friend J, Duus EM, Yan Y, Fearon KC (2016) Anamorelin in patients with non-small-cell lung cancer and cachexia (ROMANA 1 and ROMANA 2): results from two randomised, double-blind, phase 3 trials. The Lancet Oncology 17: 519–531

Tschop M, Smiley DL, Heiman ML (2000) Ghrelin induces adiposity in rodents. Nature 407: 908–13

Tschop MH, Speakman JR, Arch JR, Auwerx J, Bruning JC, Chan L, Eckel RH, Farese RV, Jr., Galgani JE, Hambly C, Herman MA, Horvath TL, Kahn BB, Kozma SC, Maratos-Flier E, Muller TD, Munzberg H, Pfluger PT, Plum L, Reitman ML et al. (2011) A guide to analysis of mouse energy metabolism. Nat Methods 9: 57–63

Tsoli M, Moore M, Burg D, Painter A, Taylor R, Lockie SH, Turner N, Warren A, Cooney G, Oldfield B, Clarke S, Robertson G (2012) Activation of thermogenesis in brown adipose tissue and dysregulated lipid metabolism associated with cancer cachexia in mice. Cancer Res 72: 4372–82

Tsoli M, Robertson G (2013) Cancer cachexia: malignant inflammation, tumorkines, and metabolic mayhem. Trends Endocrinol Metab 24: 174–83

Tsoli M, Swarbrick MM, Robertson GR (2016) Lipolytic and thermogenic depletion of adipose tissue in cancer cachexia. Semin Cell Dev Biol 54: 68–81

Tsubouchi H, Yanagi S, Miura A, Matsumoto N, Kangawa K, Nakazato M (2014) Ghrelin relieves cancer cachexia associated with the development of lung adenocarcinoma in mice. Eur J Pharmacol 743: 1–10

Vaitkus JA, Celi FS (2017) The role of adipose tissue in cancer-associated cachexia. Exp Biol Med (Maywood) 242: 473–481

van den Berg SM, van Dam AD, Rensen PC, de Winther MP, Lutgens E (2017) Immune Modulation of Brown(ing) Adipose Tissue in Obesity. Endocr Rev 38: 46–68

Vazeille C, Jouinot A, Durand JP, Neveux N, Boudou-Rouquette P, Huillard O, Alexandre J, Cynober L, Goldwasser F (2017) Relation between hypermetabolism, cachexia, and survival in cancer patients: a prospective study in 390 cancer patients before initiation of anticancer therapy. Am J Clin Nutr 105: 1139–1147

Vestergaard ET, Djurhuus CB, Gjedsted J, Nielsen S, Moller N, Holst JJ, Jorgensen JO, Schmitz O (2008) Acute effects of ghrelin administration on glucose and lipid metabolism. J Clin Endocrinol Metab 93: 438–44

Villars FO, Pietra C, Giuliano C, Lutz TA, Riediger T (2017) Oral Treatment with the Ghrelin Receptor Agonist HM01 Attenuates Cachexia in Mice Bearing Colon-26 (C26) Tumors. Int J Mol Sci 18

Wang YX, Zhu N, Zhang CJ, Wang YK, Wu HT, Li Q, Du K, Liao DF, Qin L (2019) Friend or foe: Multiple roles of adipose tissue in cancer formation and progression. J Cell Physiol

Wu J, Bostrom P, Sparks LM, Ye L, Choi JH, Giang AH, Khandekar M, Virtanen KA, Nuutila P, Schaart G, Huang K, Tu H, van Marken Lichtenbelt WD, Hoeks J, Enerback S, Schrauwen P, Spiegelman BM (2012) Beige adipocytes are a distinct type of thermogenic fat cell in mouse and human. Cell 150: 366–76

You T, Nicklas BJ (2006) Chronic inflammation: role of adipose tissue and modulation by weight loss. Curr Diabetes Rev 2: 29–37

